# Dense satellite data reveals landscape connectivity decline in the Andes-Amazon region

**DOI:** 10.1101/2022.02.17.480775

**Authors:** Paulo J. Murillo-Sandoval, Nicola Clerici, Camilo Correa-Ayram

## Abstract

There is a complex interplay of criminal groups’ control over land, illicit activities, and forest cover change in the Colombian Andes-Amazon region. This area is dealing with diverse forms of conflict and *peace,* directly impacting landscape connectivity. While many studies have documented rapid deforestation after the peace agreement in 2016, we know little about the effect of these socio-political changes on the state of landscape connectivity. We disentangle *habitat* from *connected habitat* in forest ecosystems using the Landsat archive and landscape connectivity indices. We find that in the Andes-Amazon region during 2000-2020, *connected habitat* loss reached 18%, while *habitat* loss was 13%. This result is worrisome, because it indicates that well-connected patches are more fragmented and isolated, affecting the natural connections between the Andes and Amazon biogeographical regions and the movement ability of species. The Colombian government should conduct a strategic *peacebuilding* process incorporating structural changes that prevent the increase of large-scale extractive activities that are often illegal in the region. While finding a balance between extractive activities and conservation remains a big challenge, legal land tenure, census/taxation, and specific agreements with local actors can initially prevent deforestation. We discourage localized military actions and the return of aerial fumigation of coca fields, which rather than stop deforestation might exacerbate land cover change deeper into pristine forests.

## 1. Introduction

Armed conflict is an underlying driver of human-induced landscape change. The impact of armed conflict processes on landscape evolves through periods of conflict (Hanson, 2018), which can result in human migration dynamics that increase habitat loss, forest conversion, land abandonment, and forest recovery (Barima et al., 2016; Enaruvbe et al., 2019; McNeely, 2003; Ordway, 2015). While armed conflict impacts on the landscape are diverse, conflict is a fluid process, and its effect varies according to the local institutions, actors involved, socio-ecological context and war-fighting capabilities. Although an apparent decline in human violence globally remains controversial (Cirillo & Taleb, 2016; Pinker, 2011), many countries emerge from conflict and enter into a post-conflict period. This transition from conflict to relative “*peace*” poses new socio-political and economic changes that can impact landscape change more negatively than during the conflict period (Grima & Singh, 2019).

Evidence suggests that armed conflict processes can be frequent and persistent in many countries, modifying landscape structure and connectivity (Bleyhl et al., 2017; Eklund et al., 2017; Gbanie et al., 2018; Yin et al., 2019). Although the relationship between different land cover types (e.g., forest) and conflict is well-described (Baumann & Kuemmerle, 2016), we know little about the impact of conflict processes on landscape connectivity status. Landscape connectivity is broadly defined as the degree to which the landscape facilitates or impedes species movement among resource patches (Taylor et al., 1993). Maintaining well-connected landscapes is considered an essential conservation tool to ensure evolutionary processes, ecosystem integrity, and the availability of ecosystem services that depend on organisms’ movements and their genes (Hilty et al., 2020).

The Andes Amazon Transition Belt (AATB) is a crucial location in the Neotropical realm because it allows genetic exchanges and species dispersal between the Andes and the Amazon biogeographical regions (Clerici et al., 2018). However, human pressures (Sanderson et al., 2002; Williams et al., 2020) associated with the legacy of armed conflict processes (Murillo-Sandoval et al., 2021) are threatening the dispersal capacity of species and related ecological processes, sensitive to the conversion of natural ecosystems (Haddad et al., 2015). Currently, Colombia and, most specifically, the AATB face diverse forms of conflict and *peace,* directly impacting landscape connectivity. While the 2016 peace agreement between the Government of Colombia (GoC) and the Revolutionary Armed Forces of Colombia (FARC), the largest guerrilla group, ended more than five decades of struggle, the legacy of conflict remains in formerly FARC-dissidents controlled territories. The armed conflict and FARC control over lands shaped the landscape and livelihoods and ways of economic production (Murillo-Sandoval et al., 2020). During the conflict, the AATB experienced relatively slow deforestation, given both the presence and the environmental rules created by FARC (Ruiz Serna, 2003). With the peace agreement, positive opportunities to consolidate *peace* arised for the natural environment; however, a new wave of deforestation after 2016, associated with increased land grabbing by different local and illegal actors, suggested the unforeseen consequences of this transition for landscape connectivity in the AATB region.

Earth Observation technologies offer the capacity to monitor landscape connectivity change objectively, but interpretations are often based on categorical analysis, which oversimplifies a realistic ecological assessment that involves landscape connectivity. While several connectivity approaches have been developed for different conservation tasks (Keeley et al., 2021; Saura & Pascual-Hortal, 2007), those linked with connected (reachable) and available (amount) habitats are critical for the maintenance of landscape connectivity (Saura et al., 2011a). The Equivalent Connected Area (ECA) index is a direct measurement of *connected habitat*, which considers a patch as a space where connectivity exists integrating spatial patterns and the dispersal capabilities of species (Saura & Pascual-Hortal, 2007). In this work, we disentangle forest habitat change considering three elements: *i) habitat lost* as forest habitat converted to non-natural areas (i.e., deforestation), *ii) habitat* as remaining non-transformed natural areas, and *iii) connected habitat* as non-transformed but still connected areas given a specific species dispersal distance (Fig.1SD).

Despite advances in assessing landscape connectivity dynamics (Uroy et al., 2021), tracking connectivity remains poorly addressed in conflict regions (Clerici et al., 2018; Correa Ayram et al., 2020). First, landscape connectivity change relies on simplistic descriptions of the landscape that fail when addressing the status of *connected habitat*. Therefore, disentangling *habitat lost* (e.g., deforestation) from *connected habitat* is key for practical conservation tasks (Keeley et al., 2021; Saura & Pascual-Hortal, 2007). Second, while the widespread recognition that landscapes and ecological processes are dynamic, *connected habitat* often considers a single snapshot in time (Zeller et al., 2020). It limits our ability to constantly monitor shifts in institutional frameworks (Ostrom & Nagendra, 2006); specifically in Colombia’s case: how the landscape connectivity change across different periods of armed conflict (Murillo-Sandoval et al., 2020). Finally, identifying hotspots of *connected habitat* loss is the world’s top priority to maintain ecosystem services, develop restoration campaigns and optimize resources (Tambosi & Metzger, 2013), specifically in post-conflict regions.

We overcome previous impediments by detecting intra-annual deforestation using the Continuous Change Detection and Classification algorithm (CCDC; Zhu & Woodcock, 2014) and tracking the status of *connected habitat* using the ECA index. We evaluate changes in the period 2000-2020 that encompass different stages of the Colombian conflict. This work makes two essential contributions linking landscape connectivity change in areas affected by armed conflict processes. First, we disentangle *habitat lost, habitat,* and *connected habitat* using direct measures of landscape connectivity status. Second, it recognizes the use of connectivity representations of habitat rather than static and categorical representations of landscape change. In this study, we specifically ask:

1. What was the intra-annual spatio-temporal pattern of deforestation in the Andes-Amazon region during the period 2000-2020?
2. How did *habitat* and *connected habitat* changed across different periods of the armed conflict?
3. Where are located the main hotspots of landscape connectivity loss?

## 2. Study area

Our study area is the Andes Amazon transition belt (AATB), part of the Amazon watershed (Fig 1A). It ranges from <300 m above sea level from the Amazon lowlands to 3500 m in the Eastern Andes flank. Four administrative units (departments) intersect the AATB: Putumayo, Caquetá, Guaviare, and Meta. Agricultural production, cattle ranching, and illicit coca farming are the main economic activities and the common drivers of forest conversion (Armenteras et al., 2019). Most landscape transformation occurs in its Northern region by historical human interventions, whereas the southern part remains with very low levels of change.

**Fig 1.**
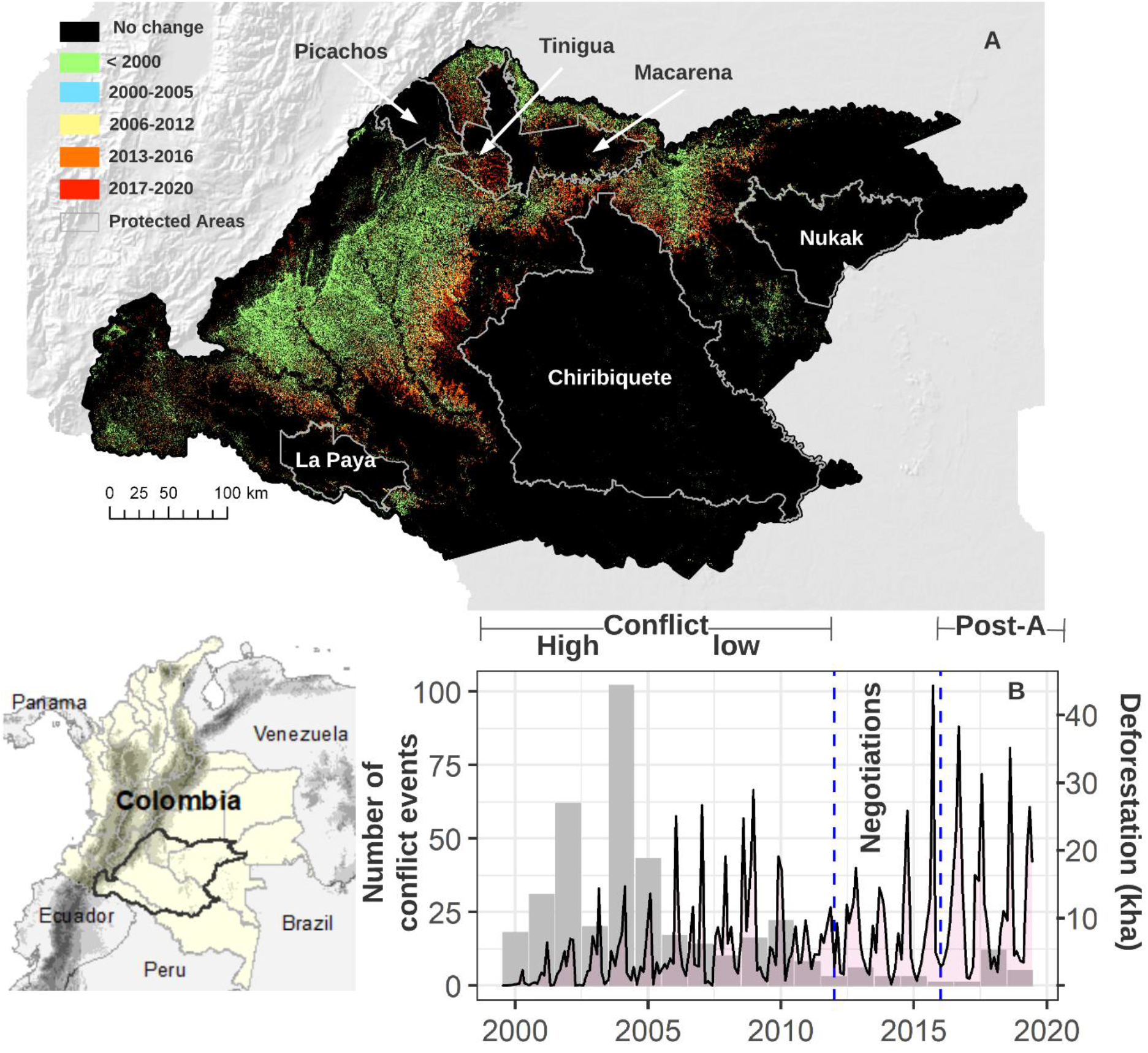
A) Deforestation accumulated for each period of conflict in the AATB; white polygons represent national protected areas. B) Intra-annual deforestation obtained from CCDC and conflict events (grey) from UCDP in different armed conflict periods.

The AATB is a biodiversity-rich area, with evidence of species dispersal between the Andes and Amazon biogeographical regions since the Miocene (~20M years ago) (Hoorn et al., 2010). However, gradual and slow habitat conversion, especially deforestation, is common in this region (Murillo-Sandoval et al., 2021). Relatively slow changes were associated with the presence of FARC that intentionally and unintentionally contributed to keeping forests saved from large-agricultural projects, the so-called ‘gunpoint conservation’ effect (Álvarez, 2003; Murillo-Sandoval et al., 2020; Rodriguez Garavito et al., 2017). The long-term control exerted by FARC produced a complex mosaic of land change (Arcila, 1989; Baptiste et al., 2017) that would seriously affect species movement and gene flow across different forest corridors (Clerici et al., 2018). The result of FARC-dissidents presence in this region since the peace agreement in 2016 led to an accelerated process of land grabbing and rapid deforestation in a desire of land consolidation and speculation.

## 3. Data and methods

Our methodology is based on two parts. First, we map intra-annual deforestation from 2000-2020 using the Continuous Change Detection and Classification algorithm (CCDC) (Zhu & Woodcock, 2014). We use the Hansen et al.(2013) tree cover map in the year 2000 and deforestation trends derived from the CCDC algorithm to derive transformed and non-transformed maps using four conflict periods: high-intensity conflict (2000-2004), low-intensity conflict (2005-2012), negotiation (2013-2016), and post-peace agreement (2017-2020) (Section 3.1). We use the definition of conflict from the Uppsala Conflict Data Program (UCDP) as: “armed force between two parties, of which at least one is the government of a state, that results in at least 25 battle-related deaths in one calendar year” (Sundberg & Melander, 2013). Our conflict periods of analysis follow Brown’s et al., (2011) progress and peace milestones. Such subdivisions involve significantly reducing conflict fatalities, cease-fire agreements, disarmament, and final peace agreement. Second, we determine the change in landscape connectivity from 2000-2020 using the ECA index and a normalized version (ECAnorm), applied to different median dispersal species distances for each conflict period (Section 3.2).

### 3.1 Transformed and non-transformed maps

CCDC is a change detection algorithm that employs all available Landsat imagery to model temporal-spectral features, including seasonality, trends, and spectral variability (Zhu & Woodcock, 2014). CCDC detects breaks into the time series that indicate the magnitude of the change, spatial location, and timing of a change. While CCDC can detect intra-annual change based on harmonic (Fourier) models, we keep the largest magnitude of change to flag a permanent change (i.e., deforestation). We further filter the magnitude of change using the Normalized Difference Fraction Index (NDFI), which is sensitive to the state of canopy forest and has provided a robust detection of deforestation across tropical forested regions (Schultz et al., 2016). We flag deforestation when magnitude values are less than zero NDFI units. We produce intra-annual deforestation maps for the period 2000-2020, following Arévalo et al., (2020) processing tools; we then reclassify the Hansen et al. (2013) tree cover dataset in 2000 as transformed (tree cover canopy < 50%) and non-transformed (tree canopy cover >50%). The non-transformed tree cover map in 2000 was updated using the deforestation events detected by CCDC for each conflict period. Finally, we carried out an accuracy assessment using 2004-2012, 2012-2016, and 2016-2020 change maps. Confusion matrices for area estimation using unbiased estimators are available in the Supplementary Data.

### 3.2 Evaluation of the landscape connectivity change

The Equivalent Connected Area (ECA) index quantified in each conflict period the *habitat* and *connected habitat* status (Saura et al., 2011b). We calculated the ECA index at two scales: 1) using the entire extent of the AATB and 2) using a hexagonal grid of 7.8 km^2^ (see Supplementary Data Fig.2SD). ECA is defined as the area of a hypothetical continuous and fully connected habitat patch, which provides the same probability of connectivity as the existing network of habitat patches in a given area (Saura et al., 2011b). The ECA index was normalized between 0-100 (ECAnorm), indicating the percentage of reachable habitat within each hexagon, and it shows how much portion of habitat is connected for species with different median dispersal distances *d* (Santini et al., 2016). We calculate ECAnorm at different median dispersal distances to cover habitat reachability for multiple species (1km-5km-10km-30km-50km-70km) (Castillo et al., 2020). We selected 10 km as a central value of the mean dispersal distance to discuss our results (Martensen et al., 2017).

At the regional level, we calculate the percentage of variation of ECA (dECA) as proposed in (Saura et al., 2011b) and compare it with the percentage of variation of habitat (dA) in each period of conflict. This comparison is convenient in terms of conservation because negative changes in *habitat* or *connected habitat* lead to a degradation of landscape’s capacity for maintaining connectivity and exchanging ecological flows. For example, suppose non-transformed areas appear distant and isolated from the previously existing ones (for the dispersal distance considered). In that case, the increase in *habitat* area will have a poor performance in terms of connectivity (dECA<dA). In contrast, if non-transformed lands present a key location within the landscape to act as connecting elements between other non-transformed lands, the increases in connectivity will be more significant than those of the transformed lands (dECA>dA). A neutral case would be one in which the entire non-transformed area belongs to the same continuous area before and after the change, in which case dECA = dA (Saura et al., 2011b).

In contrast to the regional level, we calculated dECA in each hexagon (7.8 km^2^ side) across each period of the conflict. The negative values (dECA <0) obtained were identified as connectivity losses assuming a greater intensity of loss when the values approach dECA = −100. We use the Makurhini R package (https://github.com/connectscape/Makurhini) for these analyses (Godinez y Correa, 2020).

## 4. Results

### 4.1 Intra-annual deforestation

When the Colombian armed conflict was intense, deforestation recorded was slower, whereas the beginning of the peace agreement led to rapid deforestation. Annual deforestation increases for each change period. In 2004-2012 deforestation was 56555±13500ha, in 2012-2016, 71283±25574ha, and after the peace agreement (2016-2020) 95300 ± 8894ha (Fig 1A). These trends indicate that an increase in deforestation rates accompanies the de-escalation of conflict. Lower deforestation area variability coincides with more conflict events from the Uppsala database (Sundberg & Melander, 2013) during the conflict period (Fig 1A).

In contrast, a decrease in conflict events denotes more deforestation during the low-conflict and negotiations periods (Fig 2A). The uncertainty associated with the negotiation process increased deforestation, whereas the consolidation of the agreement in 2016 almost doubled deforestation immediately in the region. These findings have been documented in previous studies (Murillo-Sandoval et al., 2020). Still, our detailed long-term analysis shows the complexity of the armed conflict process leading to specific repercussions in deforestation trends. For instance, the widespread expansion of illegal land markets exacerbating land grabbing in new locations linked with a repressed search for access to land that was not possible when FARC exerted control over the region.

**Fig 2.**
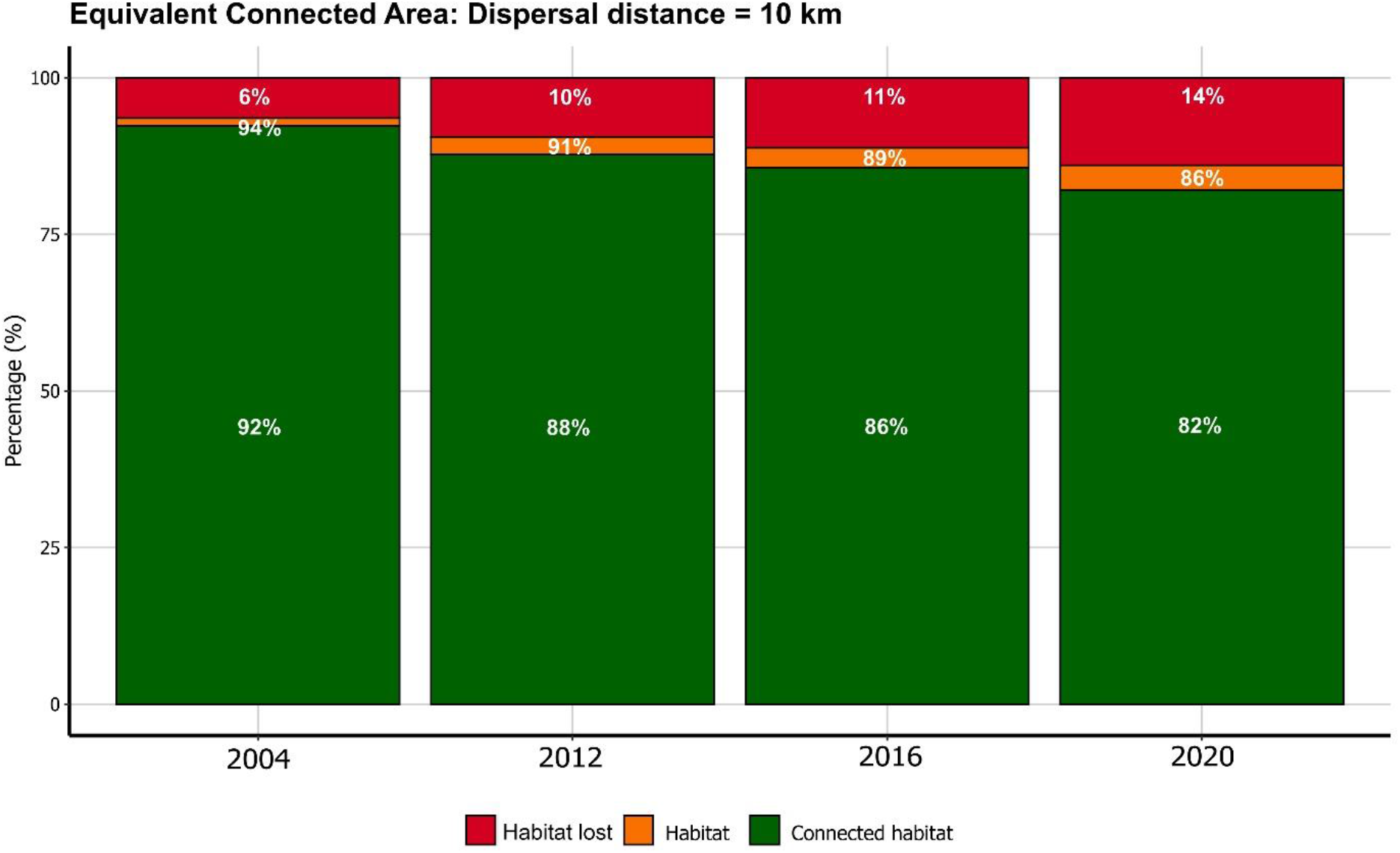
Distribution (%) of habitat lost, habitat, and connected habitat for the AATB.

### 4.2 Changes in landscape connectivity

We found that the loss of *connected habitat* has been considerably greater than the loss of *habitat*. By 2020, the AATB region lost 13% of *habitat*, and 18% of *connected habitat* present initially in the region. In the change period 2004-2020, 2 Mha (11%) of *connected habitat* and 1.4 Mha (8%) of *habitat* were lost. We find a similar trend during each of the armed conflict periods (Fig 2). The analysis of dECA and dA for each conflict period suggests a quick loss of *habitat* and *connected habitat*. During low-conflict periods centered in 2004 and 2012 (8 years) the dECA value is −5% and dA = −2.4%. In the negotiation period 2012-2016 (4 years) the loss of connectivity is dECA = −2.4% and habitat achieved dA = −1.8 %. In contrast, during the post-agreement period (4 years), the loss of connectivity and habitat increased again by dECA= −4% and dA= −3%, respectively.

Temporal distribution of connectivity shows a detrimental effect in the ECAnorm values at hexagon-level. ECAnorm values changed from 91% to 83% between 2004 and 2020, respectively. This trend is evident from the western Andean foothills to the Amazonian lowlands of Sabanas del Yari, Chiribiquete, the south of Guaviare, Tinigua, and the Chiribiquete-Macarena corridor (Fig 3C), see ECAnorm for each period in Supplementary Data (Fig.3SD). As expected, ECAnorm values also decrease once dispersal distance capabilities increase (d>10 km) (see Supplementary Data Fig4.SD). We also identified areas with stable connectivity associated with riparian forests surrounding the Caquetá and Caguán rivers (Fig 3A, 3B).

**Fig 3.**
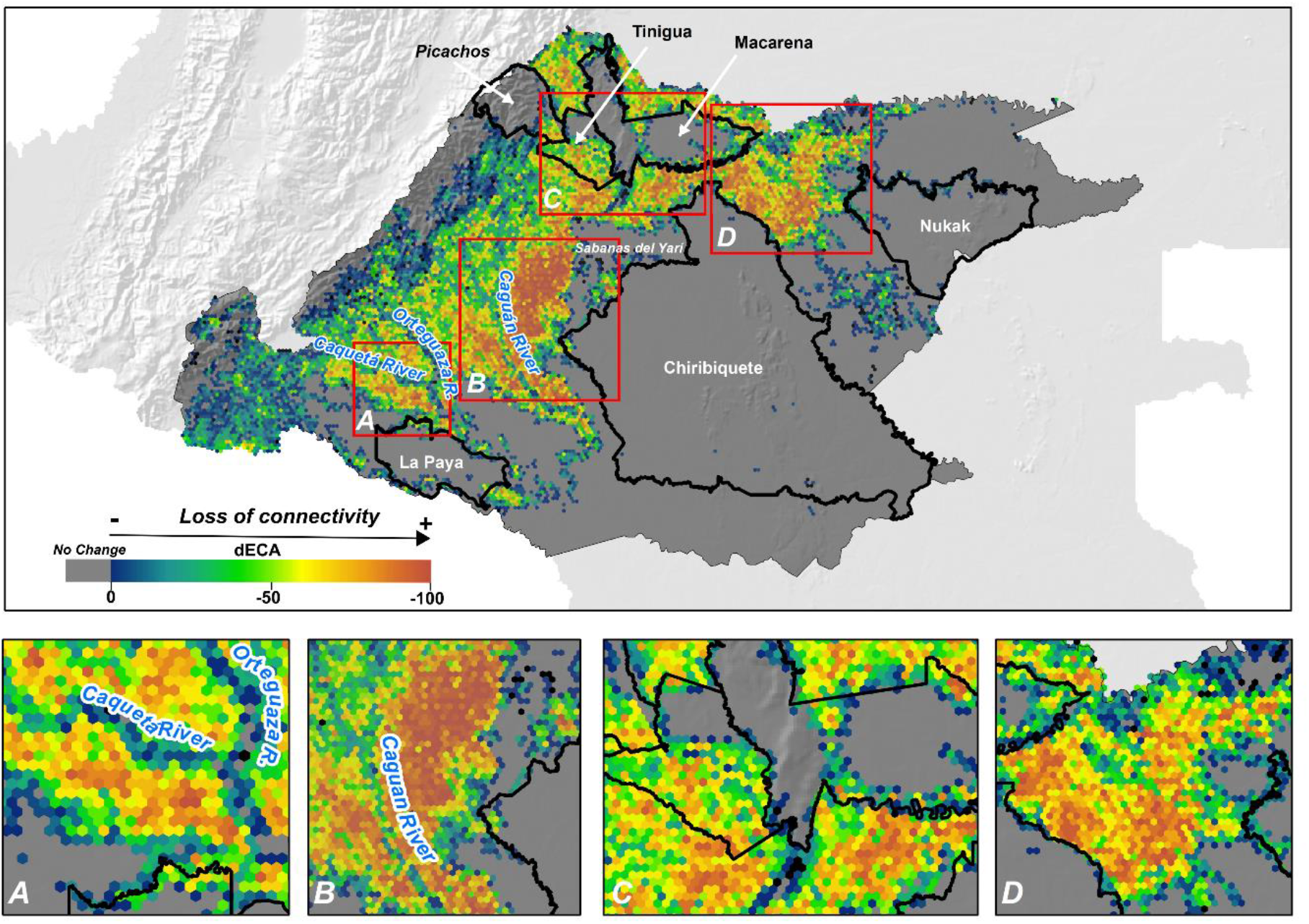
Hotspots of loss in dECA at hexagon-level for the period 2004-2020. A) Puerto Guzman, B) Chiribiquete (western) C) the corridor Tinigua-Macarena and D) the corridor Macarena-Nukak.

### 4.3 Hotspots of loss in connected habitat

We identify four local regions with high rates of connectivity loss: Puerto Guzman, Chiribiquete (western), the corridor Tinigua-Macarena and the corridor Macarena-Nukak (Fig 3). The first hotspot in Puerto Guzman is very near to the La Paya National Natural Park. *Connected habitat* loss expands to the southeast of AATB, forming a disturbed region that disconnects La Paya and Chiribiquete. The hotspot western to Chiribiquete covers the park’s limit, the Sabanas del Yari and Caguan river, increasing pressure over remaining forest relics between the Andean foothills and Amazonian lowlands. The case of the Tinigua-Macarena corridor (Fig 3C) is shocking. Losing *connected habitat* in this corridor indicates a growing division with Chiribiquete (north), disconnecting a key bridge for gene flow interchange between the Andes and Amazon regions (Clerici et al., 2018). Finally, the corridor Macarena-Nukak coincides with the expansion of transformed lands in the Guaviare department (Fig 3D). The rapid deforestation in this region affects species richness and mobility, and indigenous communities that are more exposed to land conflicts with farmers and criminal groups for cattle ranching and coca farming production.

## 5. Discussion

Our findings are alarming because we found that in the AATB the loss of *connected habitat* is considerably higher than the loss in *habitat*. The loss of critical remaining habitat patches harms the movement ability of species with limited dispersal capacity (Herrera et al., 2017), mining metapopulations resilience, and persistence. Such regional connection is essential for species dispersal, supporting genetic exchange and evolutionary processes through the diversity of Neotropical ecosystems. Phylogenetic evidence showed the importance of this highland–lowland connectivity for the foundation and maintenance of regional mammals, avian and invertebrate biodiversity (Elias et al., 2009; Haag et al., 2007; Hines et al., 2011; McGuire et al., 2014). Our findings indicate that hotspots of connectivity loss are also located inside Protected Areas (e.g., Tinigua), where conservation aims to protect ecosystems of outstanding ecological and scientific value.

The consequences of the peace agreement in *connected habitat* are shocking. We find that the loss in *connected habitat* during the low-conflict period (8 years) was similar to that observed during only the post-peace agreement period (4 years). The widespread loss of *connected habitat* after the peace agreement indicates that the reachable habitat for species is now less available given the quick transformation of large forest areas with high internal connectivity. While it is common to anticipate an increase in deforestation in the transition conflict to *peace* (Grima & Singh, 2019), it is less common to observe a rapid disconnection between forest relics after a few years of socio-political transitions. The rise in connectivity loss associated with the exit of FARC drove a shift in the institutional framework, which might threaten the future provision of services in the AATB due to loss of ecological integrity (Salazar et al., 2018).

In vast forested areas, such as in the Andes-Amazon region, conservation aims to protect the entire natural landscape, ideally by working with indigenous and local farmer communities (Keeley et al., 2021). However, controlling *connected habitats* in regions affected by a long-lasting armed conflict requires an extra effort between different global, national and local agencies to ensure ecosystem processes maintenance. *Connected habitat* helps to develop zero-deforestation initiatives that currently proliferate in Colombia (Furumo & Lambin, 2020) and Sustainable Land-use systems as alternatives to conventional unproductive cattle ranching (Bonatti et al., 2021). The extreme loss of *connected habitat* observed indicates that the application of these initiatives remained very weak as tools for effective environmental conservation and governance (Furumo & Lambin, 2020; Krause, 2020). The structural reasons for deforestation are not taken deeply into account through these initiatives or other national policies. While zero-deforestation initiatives claim to preserve the environment, the Colombian government still supports an extractive model that poorly addresses local demands for legal land tenure and infrastructural investment. This discrepancy is also motivated for future private interests shaping the connection region between the Andes and Amazon biomes.

This study illustrates that management actions to preserve *connected habitat* depend not only on the amount of habitat available but also on how accessible it is for the species. The division of AATB in hexagons allows identifying the sites with the lowest to the highest loss of connectivity per unit area, providing a more detailed view of the anthropic matrix consolidation and where conservation decisions should be on track. In sites with a high amount of habitat and well connected (e.g., areas of no change in grey, Fig 3) for species with different dispersal capacities, conservation efforts should be directed to maintain what remains, primarily through the implementation of Protected Areas. However, sites that have lost more than 50% of their *connected habitat* (e.g., yellow to red hexagons in Fig 3) could be classified as a high priority for restoration (Tambosi & Metzger, 2013). Restoration strategies aimed at increasing the size of patches (> amount of habitat) and at reconnecting habitat through the implementation of corridors (>connected habitat).

Some limitations need to be mentioned. First, while using Landsat, we detected deforestation at 0.27ha (3 pixels); our connectivity assessment considers landscapes with a minimum mapping unit of 10 ha. Consequently, we cannot fully capture very fine-scale patterns, such as small patches or remnants of not transformed areas suitable for species with limited dispersal abilities, i.e., acting as stepping stones (Saura et al., 2011a). Second, we do not consider data on dispersal abilities and other relevant ecological information of specific species. However, we employed a broad threshold of median dispersal distances that is widely representative of a group of species that identifies in detail where connectivity changes (Castillo et al., 2020; Gonzalez-Borrajo et al., 2017). Third, habitat change was characterized by armed conflict periods; however, regional causal mechanisms were not addressed for each hotspot of change detected. While causal mechanisms require fieldwork, the vast literature associated with armed conflict and forest coincides with specific socio-political changes occurring during the last five years (Mendoza, 2020; Murillo-Sandoval et al., 2021; Reilly & Parra-Peña, 2019).

Future work should focus on national-level connectivity analysis and incorporate the effect of human pressures in evaluating landscape connectivity change (see dECA_matrix_ in Saura et al., 2011a). For example, integrating landscape resistance to dispersal through a legacy-adjusted human footprint index (Correa Ayram et al., 2020), as reported in previous studies at continental (Barnett & Belote, 2021; Belote et al., 2017) and ecoregional level (Castillo et al., 2020). Several regions in Colombia require further investigation incorporating prospective regional scenarios (Gonzalez-Borrajo et al., 2017) for planning future conservation efforts and goals (González-González et al., 2021).

## 6. Conclusions

We use dense satellite data to unveil landscape connectivity change in the AATB during the last 20 years. While deforestation increased drastically in the post-peace agreement period, its magnitude is less critical than the massive loss in *connected habitat,* suggesting the rapid fragmentation of remaining well-connected forest patches. The peace agreement established an initial route to building sustainable development with local communities. However, a comprehensive understanding of local communities’ needs and long-term projects that benefits them remains poorly implemented. The Colombian government should focus on implementing stable governance structures and strategies to work with local farmers and indigenous populations, rather than localized military actions and the aerial fumigation of coca, which are not effective mechanisms that could halt *habitat connectivity* loss.

