## Supplementary for "Dense satellite data reveals landscape connectivity decline in the Andes-Amazon region"

### Graphical definitions for habitat lost, habitat and connected habitat

A simple case to illustrate the different types of landscape elements related to the evaluation of connectivity change in the AATB. The *habitat lost*, shown in clear green referred as transformed lands that originally were habitat. *Habitat*, shown in dark green as non-transformed lands that can appear as isolated patches. *Connected habitat*, as a set of fully connected habitat patches given a considered dispersal distance (d). In this illustrative example, d= 10 km.

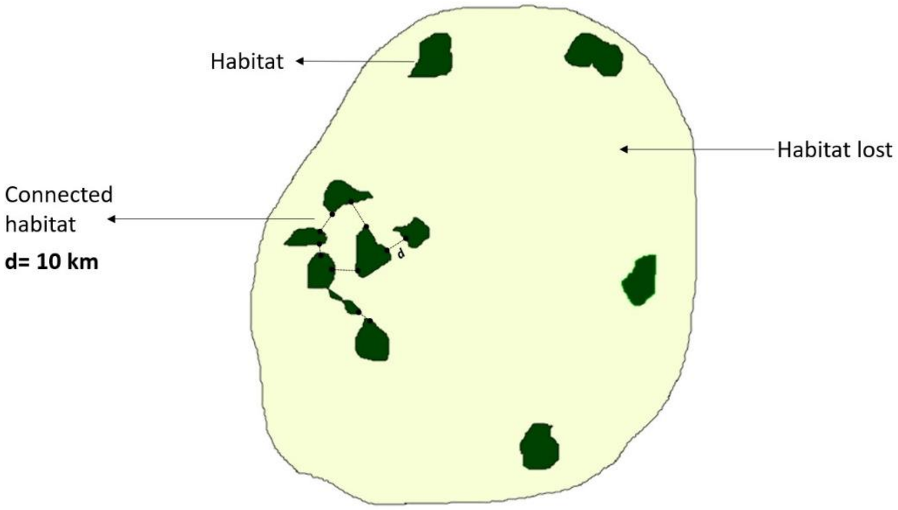


**Fig 1SD.** Scheme to explain different types of landscape elements

### Area estimation

A random stratified procedure was created to evaluate areas estimation. We set up a target error of 0.02 and overall accuracies for four classes as follows:

1. non-transformed (stable Forest) 80%
2. non-transformed to transformed (disturbances) 70%
3. Transformed (stable non-forest) 80% and
4. Buffer non-transformed stratum surrounding non-transformed to transformed (80%)

We distribute random independent samples. A total of 430 for each change map (2004-2012, 2012-2016, and 2016-2020) as follows:

1. non-transformed (stable Forest): 300
2. non-transformed to transformed (disturbances): 30
3. Transformed (stable non-forest): 50
4. Buffer non-transformed stratum surrounding non-transformed to transformed: 50

**Confusion matrices and accuracies**

Columns = Reference Labels, Rows = Map Values.

Pixel-based confusion matrix 2004-2012

| Class | 1 | 2 | 3 | 4 |
|---|---|---|---|---|
| 1 | 299 | 0 | 1 | 0 |
| 2 | 4 | 46 | 0 | 0 |
| 3 | 1 | 0 | 49 | 0 |
| 4 | 29 | 1 | 0 | 0 |

Area proportions confusion matrix 2004-2012

| Class | 1 | 2 | 3 | 4 |
|---|---|---|---|---|
| 1 | 0.818 | 0 | 0.003 | 0 |
| 2 | 0.002 | 0.02 | 0 | 0 |
| 3 | 0.002 | 0 | 0.074 | 0 |
| 4 | 0.078 | 0.003 | 0 | 0 |

Accuracies 2004-2012

| Property/Strata | 1 | 2 | 3 | 4 |
|---|---|---|---|---|
| Area proportion | 0.9 | 0.023 | 0.077 | 0 |
| Standard Error (area prop) | 0.004 | 0.003 | 0.003 | 0 |
| Area (ha) | 17585644 | 452446 | 1502418.2 | 0 |
| 95% CI (ha) | 161670 | 108583.9 | 119777.86 | 0 |
| Producer's | 0.909 | 0.883 | 0.964 | 0 |
| User's | 0.997 | 0.92 | 0.98 | 0 |

Overall accuracy 0.913 ± 0.006

Pixel-based confusion matrix 2012-2016

| Class | 1 | 2 | 3 | 4 |
|---|---|---|---|---|
| 1 | 299 | 0 | 1 | 0 |
| 2 | 7 | 43 | 0 | 0 |
| 3 | 0 | 1 | 49 | 0 |
| 4 | 29 | 1 | 0 | 0 |

Area proportions confusion matrix 2012-2016

| Class | 1 | 2 | 3 | 4 |
|---|---|---|---|---|
| 1 | 0.835 | 0 | 0.003 | 0 |
| 2 | 0.002 | 0.011 | 0 | 0 |
| 3 | 0 | 0.002 | 0.096 | 0 |
| 4 | 0.049 | 0.002 | 0 | 0 |

Accuracies 2012-2016

| | 1 | 2 | 3 | 4 |
|---|---|---|---|---|
| Area proportion | 0.887 | 0.015 | 0.099 | 0 |
| Standard Error (area prop) | 0.003 | 0.003 | 0.003 | 0 |
| Area (ha) | 17326105 | 285133.4 | 1929270.1 | 0 |
| 95% CI (ha) | 127650.7 | 102298.3 | 130674.39 | 0 |
| Producer's | 0.942 | 0.749 | 0.972 | 0 |
| User's | 0.997 | 0.86 | 0.98 | 0 |

Overall accuracy 0.942 ± 0.007

Pixel-based confusion matrix 2016-2020

| Class | 1 | 2 | 3 | 4 |
|---|---|---|---|---|
| 1 | 300 | 0 | 0 | 0 |
| 2 | 5 | 45 | 0 | 0 |
| 3 | 0 | 0 | 50 | 0 |
| 4 | 30 | 0 | 0 | 0 |

Area proportions confusion matrix 2016-2020

| Class | 1 | 2 | 3 | 4 |
|---|---|---|---|---|
| 1 | 0.794 | 0 | 0 | 0 |
| 2 | 0.002 | 0.02 | 0 | 0 |
| 3 | 0 | 0 | 0.111 | 0 |
| 4 | 0.074 | 0 | 0 | 0 |

Accuracies 2012-2016

| | 1 | 2 | 3 | 4 |
|---|---|---|---|---|
| Area proportion | 0.87 | 0.02 | 0.111 | 0 |

| | | | | |
|---|---|---|---|---|
| Standard Error (area prop) | 0.001 | 0.001 | 0 | 0 |
| Area (ha) | 16998035 | 381202 | 2161271.4 | 0 |
| 95% CI (ha) | 35578.85 | 35578.85 | 0 | 0 |
| Producer's | 0.913 | 1 | 1 | 0 |
| User's | 1 | 0.9 | 1 | 0 |

Overall accuracy 0.924 ± 0.002

**Calculating grid hexagon size**

We use the Shannon diversity index (SHDI) to calculate the optimal grid size unit that captures the diversity of the connected landscape. We employ the transformed non-transformed map in 2020 to calculate SHDI spanning from 50 m to 4000 m surrounding a set of 500 points randomly sampled across the study area. We identify the second instance under which the curve became flat.  This location is known as the “elbow” or “knee” where the curve stops increasing. We calculated the Unit Innvariant Knee (UIK) for each spatial incremental buffer and calculated the mean value after removing values that indicated no data and where buffers contain more than 80% of land change information. We obtained a value of 2.22 km that was used as the main radio for hexagon generation leading a hexagon area of 7.8 km2. Calculations were made using an inflection package available in R (Christopoulos, 2016).

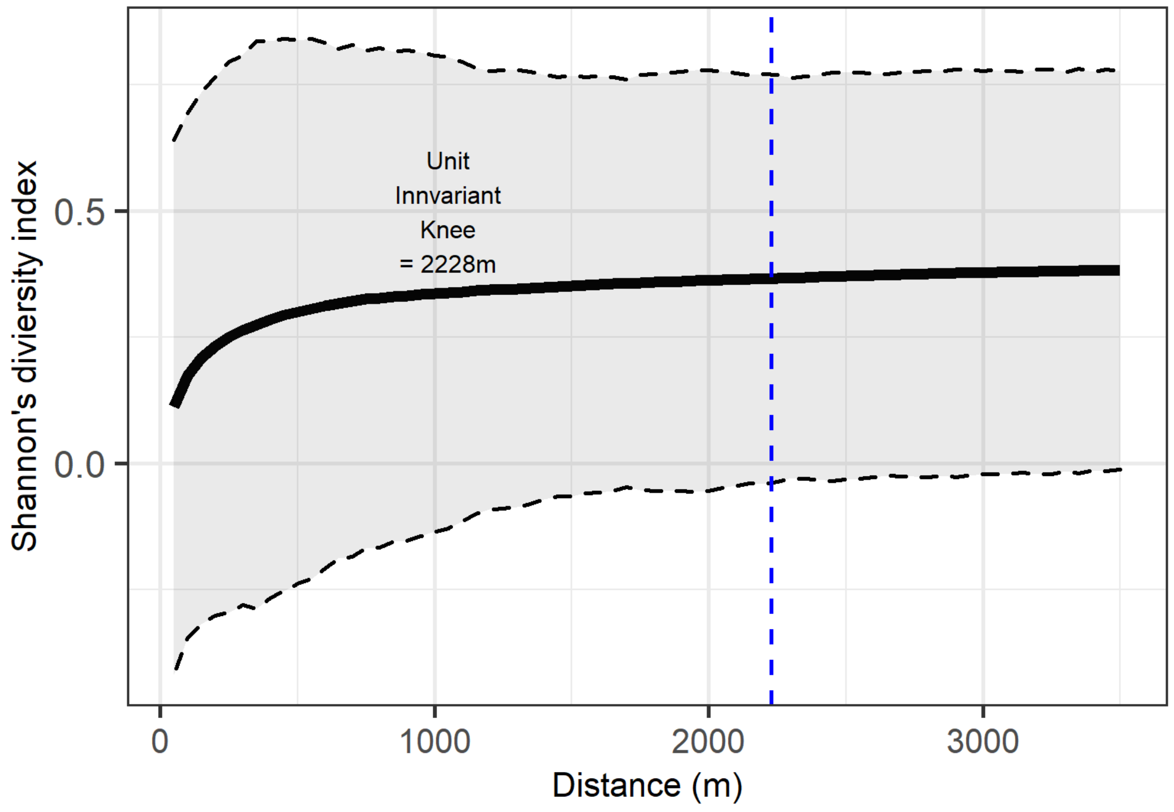

**Fig 2SD**. Shannon's diversity index across random points in the AATB. The black line denotes the mean value across all buffers conflict events. The dashed line represents the variations of individual random points. The blue line identifies the UIK value.

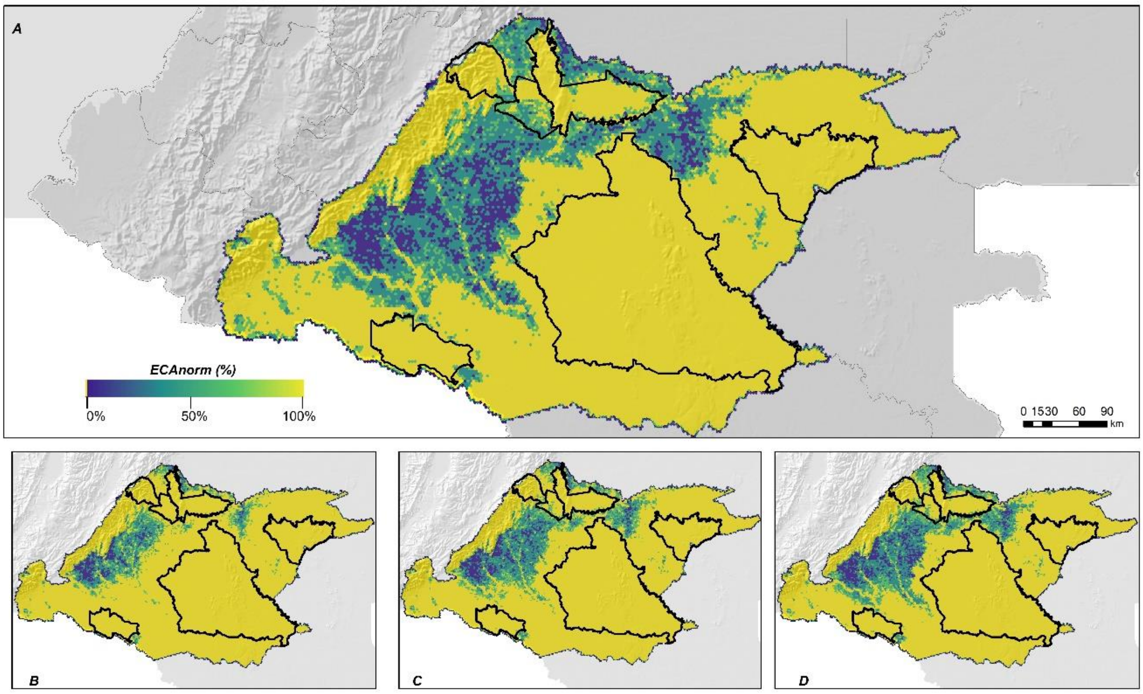


**Fig 3SD.** ECAnorm for the period 2000-2020 (A). ECAnorm for each conflict period 2004-2012 (B), 2012-2016 (C) and 2016-2020 (D).

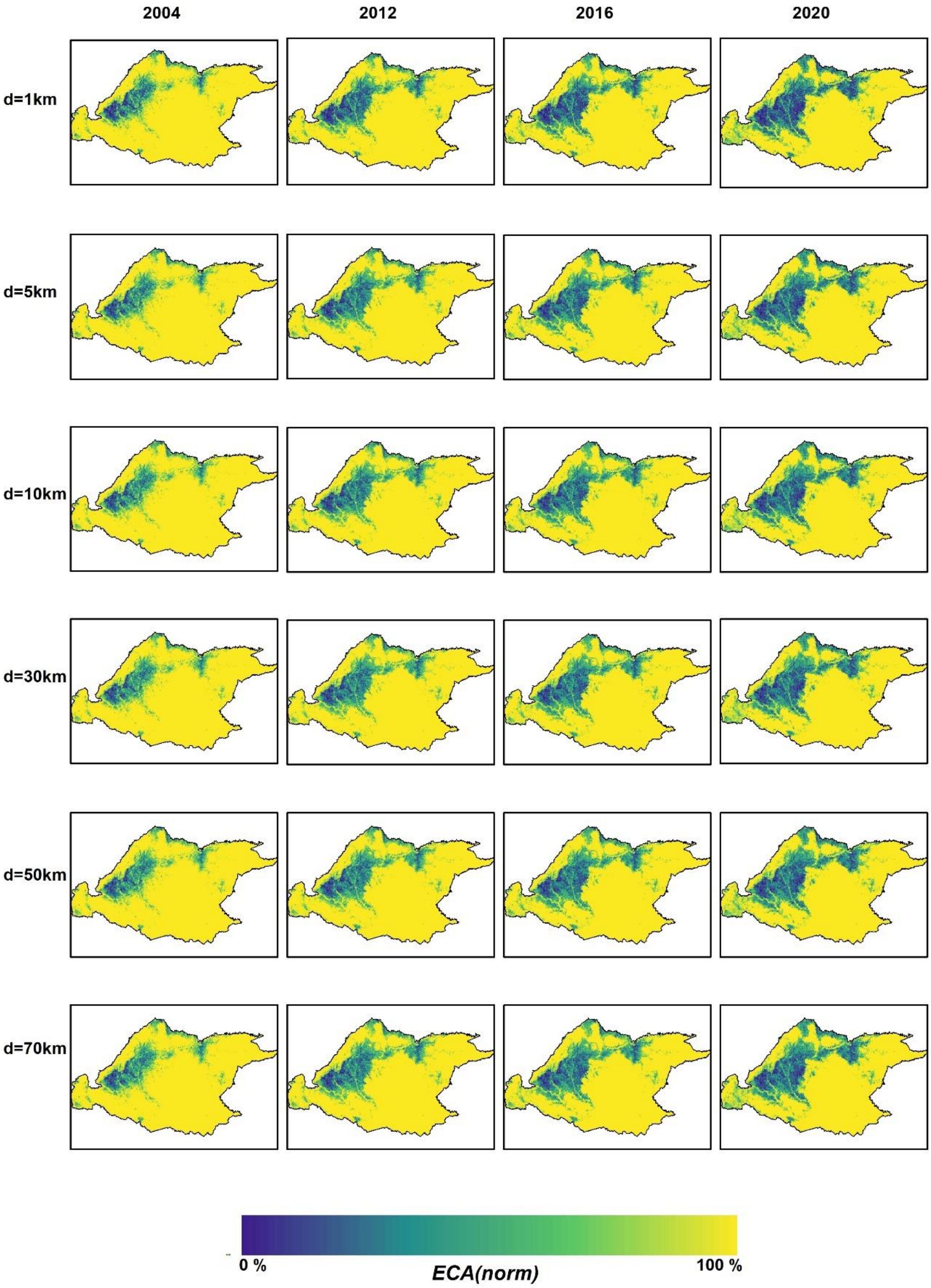


**Fig 4SD**. ECAnorm for each period of analysis and for different dispersal distances.
